# Liver kinase B1 (LKB1) regulates the epigenetic landscape of mouse pancreatic beta cells

**DOI:** 10.1101/2024.05.13.593867

**Authors:** Nejc Haberman, Rebecca Cheung, Grazia Pizza, Nevena Cvetesic, Dorka Nagy, Hannah Maude, Lorea Blazquez, Boris Lenhard, Inês Cebola, Guy A. Rutter, Aida Martinez-Sanchez

**Affiliations:** MRC London Institute of Medical Sciences, London W12 0NN, UK; Institute of Clinical Sciences, Faculty of Medicine, Imperial College London, London W12 0NN, UK; Division of Neuroscience, Department of Brain Sciences, Imperial College London, London, UK; Section of Cell Biology and Functional Genomics, Faculty of Medicine, Imperial College London, London, UK; Section of Genetics and Genomics, Department of Metabolism, Digestion and Reproduction, Imperial College London, London, UK; National Heart and Lung Institute, Imperial College London, London, UK; Department of Neurosciences, Biogipuzkoa Health Research Institute, 20014 San Sebastián, Spain; Ikerbasque, Basque Foundation for Science, 48009 Bilbao, Spain; CIBERNED, ISCIII (CIBER, Carlos III Institute, Spanish Ministry of Sciences and Innovation), 28031, Madrid, Spain; Research Centre of the Centre Hospitalier de l’Université de Montréal (CRCHUM), Faculté de Médecine, Université de Montréal, Montréal, QC, Canada; Lee Kong Chian Medical School, Nanyang Technological University, Singapore

## Abstract

Liver kinase B1 (LKB1/STK11) is an important regulator of pancreatic β-cell identity and function. Elimination of *Lkb1* from the β-cell results in improved glucose-stimulated insulin secretion and is accompanied by profound changes in gene expression, including the upregulation of several neuronal genes. The mechanisms through which LKB1 controls gene expression are, at present, poorly understood. Here, we explore the impact of β cell-selective deletion of *Lkb1* on chromatin accessibility in mouse pancreatic islets. To characterize the role of LKB1 in the regulation of gene expression at the transcriptional level, we combine these data with a map of islet active transcription start sites and histone marks. We demonstrate that LKB1 elimination from β-cells results in widespread changes in chromatin accessibility, correlating with changes in transcript levels. Changes occurred in hundreds of promoter and enhancer regions, many of which were close to neuronal genes. We reveal that dysregulated enhancers are enriched in binding motifs for transcription factors important for β-cell identity, such as FOXA, MAFA or RFX6 and we identify microRNAs (miRNAs) that are regulated by LKB1 at the transcriptional level. Overall, our study provides important new insights into the epigenetic mechanisms by which LKB1 regulates β-cell identity and function.

## INTRODUCTION

Adequate insulin secretion is required for the maintenance of normal glucose homeostasis in mammals. Insulin is stored and released from the β-cells which comprise 60-70 % of pancreatic islet cells(1). Deficiencies in this process are ultimately responsible for all forms of diabetes mellitus(2), a disease which affects more than 1 in 10 adults worldwide (https://diabetesatlas.org/atlas-reports/).

The highly conserved Ser/Thr protein kinase Liver Kinase B1 (LKB1/STK11) is a tumour suppressor that restricts cell growth in mammalian cells. Loss-of-function mutations in LKB1 cause Peutz-Jegers syndrome, a familial disorder characterized by the development of spontaneous intestinal tumours (3) (4).

LKB1 is also a master regulator of pancreatic β-cell size, polarity and function (5–7). Recently, LKB1 has also emerged as a target for glucagon-like peptide-1 receptor (GLP-1R) agonists in the β-cell (8). Agonism at these receptors, both by the endogenously secreted incretin hormones and by pharmacologically-administered drugs, is an important means of regulating insulin secretion in health and in type 2 diabetes (T2D) (9, 10).

LKB1 functions *via* the activation of several downstream kinases from the AMP-activated protein kinase (AMPK) family (11). It was initially proposed (12, 13) that LKB1 function in β- cells was mostly mediated by AMPK itself. However, it was later demonstrated that LKB1 and AMPK have only partially overlapping functions in this cell type(6) with deletion of AMPK inhibiting(14), whilst that of LKB1 activating (7) insulin secretion. Since improved insulin secretion persists in the long term (>10 months) in mice with β-cell specific LKB1 ablation (LKB1βKO) (5), mimicking this pro-secretory activity of LKB1βKO with drugs that target this, or downstream enzymes, represents an attractive approach for the treatment of diabetes. However, given the broad impact of inhibiting LKB1 across multiple tissues (15), further work is required to understand the precise molecular mechanisms underlying increased insulin secretion after LKB1 deletion.

Interestingly, increased glucose-stimulated insulin secretion (GSIS) upon LKB1 inactivation occurs despite a strong and progressive deterioration of mitochondrial ultrastructure and function over time, as evident by electron microscopy (16). Other changes included altered cellular morphology and polarity, without major alterations in β-cell mass (7, 13, 14). We have previously shown that broad changes in gene expression in LKB1βKO islets are likely to underlie the functional and morphological alterations observed (5, 6). Thus, LKB1βKO islets displayed altered expression of almost 3000 genes, with a strong enrichment for those involved in neuronal pathways as well as increased expression of hepatic, tumoral, glutamate signalling and α-cell specific genes (6). Increased glutamate signalling could contribute to the enhanced insulin secretion of LKB1βKO β-cells (5, 6). Interestingly, some, but not all, of these changes overlapped with those occurring in islets with deletion of AMPK catalytic subunits (AMPKβdKO) in β-cells using the same Cre deleter strain (6). Whilst we have previously shown that dysregulation of microRNA (miRNA) expression likely contributes to the loss of cellular identity in AMPKβdKO islets, whether these important regulators of gene expression and of cellular estates (17) also contribute to loss of cellular identity in LKB1βKO β-cells remains to be interrogated.

LKB1 has also been shown to maintain cellular identity in other systems. For example, in Treg cells (18) LKB1 regulates cell identity *via* activation of TGFβ signalling-Smad2/3 transcriptional activity and through changes in DNA methylation of specific *loci*. Similarly, LKB1 drives reprogramming from progenitor-like to alveolar type II-like states in lung cancer(19). Mechanistically, LKB1 has been shown recently to control metastatic transformation of lung adenocarcinomas by promoting changes in chromatin accessibility and transcriptional regulation(20). Neither the extent to which transcriptional regulation contributes to LKB1-dependent changes in gene expression in islet cells, nor the underlying mechanisms, have been investigated.

Active *cis*-regulatory elements are central to transcriptional regulation and are characterised by “open” chromatin that is amenable to enzymatic digestion, reflecting the displacement of nucleosomes by transcription factors and other components of the transcriptional machinery (21). With this in mind, here, we performed Assay for Transposase-Accessible Chromatin with high-throughput sequencing (ATAC-seq) to profile active *cis*-regulatory elements in islets from LKB1βKO and control animals. We used Super-Low Input Carrier Cap Analysis of Gene Expression (SLIC-CAGE)-identified transcription start sites (TSS) and existing genome-wide maps of islet histone marks to further characterize the role of transcriptional regulation in LKB1-mediated control of gene expression in mouse β-cells.

We show that LKB1 elimination has a widespread impact on chromatin accessibility, with hundreds of enhancer and promoter regions displaying increased accessibility. These dysregulated regions are enriched in binding motifs for important islet transcription factors required for normal β-cell function, as well as widely-expressed transcription factors involved in cell survival and proliferation. Finally, we integrated islet ATAC-seq and SLIC-CAGE data to generate a precise, islet-specific map of microRNA (miRNA) TSS and to identify miRNAs regulated by LKB1 at the transcriptional level.

Our findings provide important insights into the molecular mechanisms by which LKB1 controls β-cell identity and function and identify LKB1 as a major modulator of the epigenetic landscape of the β-cell.

## MATERIALS AND METHODS

### Mice maintenance

LKB1βKO (LKB1^fl/fl^, Ins1-Cre^+/–^) and control (LKB1^fl/fl^, Ins1-Cre^−/−^) mice(6) were housed with free access to standard mouse chow diet at the Imperial College Central Biomedical Service and approved by the College’s Animal Welfare and Ethical Review Body according to the UK Home Office Animals Scientific Procedures Act, 1986 (Project Licenses PA03F7F0F and PP7151519).

### Isolation and maintenance of islets

Mouse islets were isolated using collagenase (Roche) digestion as previously described(22). Islets were allowed to recover overnight preceding SLIC-CAGE and miRNA RT-qPCR experiments in RPMI 1640, 10% fetal bovine serum (FBS), L-glutamine, and 11 mM glucose at 37°C with 95% O2/ 5% CO2.

### RNA extraction and miRNA reverse transcription and quantitative PCR (RT-qPCR)

RNA was extracted using Trizol (Thermo Fisher Scientific) following the manufacturer’s instructions. MiRNA reverse transcription (RT) and qPCR was performed from 30ng RNA as previously described(23) with miRCURY LNA probes and miRCURY LNA RT and SYBR® Green PCR Kits. Let-7d-3p and miR-573-3p (data not shown) were used as endogenous controls due to their stable expression in our system(23).

### ATAC-seq and SLIC-CAGE

ATAC-seq was performed as previously described(24, 25) using 100 hand-picked islets per mouse (4 littermate LKB1βKO and 4 control female mice, 14 weeks old). Briefly, isolated islets were collected in ice-cold phosphate buffer saline (PBS) and pelleted at 100 rcf at 4°C for 1 min. Nuclei were isolated by resuspending the islets in 300 µl of cold ATAC lysis buffer (10 mM Tris-HCl, pH 7.4, 10 mM NaCl, 3 mM MgCl2 and 0.1% IGEPAL CA-630). After a 20 min incubation on ice with gentle shaking, the nuclei were released by passing the lysate through a 1ml syringe (with needle attached) 20 times. Nuclei were pelleted at 500 rcf at 4°C for 10 min, then washed once by resuspending in 100 µl of cold ATAC lysis buffer. At this point, nuclei were counted in a TC20 Automated Cell Counter (Bio-Rad). 50,000 nuclei were then transferred to a new 1.5 mL Eppenforf tube and pelleted as previously. The transposition reaction was carried out immediately afterwards using 25 µL 2x TD buffer (Illumina, Nextera DNA Library Preparation), 2.5 µL transposase (Illumina, Nextera DNA Library Preparation), 22.5 µL nuclease-free water. The transposition mix was incubated at 37°C for 30 min in a thermomixer with constant shaking (1,000 rpm). The reaction was then immediately purified using Qiagen MinElute Reaction Cleanup Kit, following manufacturer’s instructions (Qiagen, 28004) and eluted in 21 µL of Elution Buffer. ATAC libraries were generated by initial amplification for 5 cycles using the following PCR conditions: 72°C 5 min; 98°C 30 s; then cycling at 98°C 10 s, 63°C 30 s and 72°C 1 min using 25 µL NEBNext High-Fidelity 2x Master Mix, 2.5 µL of 25 µM Forward/Reverse ATAC-seq index primers(26) and 20 µL of the purified transposed DNA. The number of additional PCR cycles needed to reach sufficient library amplification was estimated as previously described(26). Amplified libraries were purified using Qiagen MinElute PCR Purification Kit (Qiagen, 28004), eluted in 20 µL of nuclease-free water, and quantified using the KAPA Library Quantification kit for ABI Prism (Roche, KK4854). ATAC libraries were pooled at equimolar ratios prior to sequencing in a HiSeq4000 instrument with paired-end 50bp reads at the Genomics Unit of the Centre for Genomic Regulation (Barcelona, Spain).

SLIC-CAGE was performed as in (27) from 100 ng of total RNA extracted as described above from islets isolated from a 18 week old female mouse on a C57BL6/J background. 12 cycles were performed for library amplification. The amplified library was purified with AMPure XP beads and visualised using a HS DNA chip (Bioanalyzer, Agilent). The library was sequenced on a HiSeq2500 instrument with paired-end, 150bp reads at the Genomics Facility, MRC, LMS).

### ATAC-seq and ChIP-seq data analysis

Adaptors were trimmed from the ATAC-seq reads using Trim Galore v0.4.1 (https://github.com/FelixKrueger/TrimGalore) with the –quality 15 parameters for paired end data and reads aligned to mm10 with Bowtie2 v2.4.4(28) with the parameters -p 12 -- no-unal. Picard v2.23.3 (http://broadinstitute.github.io/picard/) “CollectInsertSizeMetrics” was used to assess library size distribution. Bowtie2 v2.4.4 was also used for the alignment of ChIP-seq reads, downloaded from Lu et al. (29) (GSE110648, H3K4me3, H3K4me1, inputs) and Urizar et al.(30) (GSM3012378, H3K27ac, input) with the parameters -p 12 --no-unal, for paired end and single end data, respectively. MACS2(31) v2.1.0 was used for peak calling with the following parameters: -f BAMPE --bw=300 -B -q 0.01 --nomodel --shift 37 --extsize 73 for ATAC-seq and -f BAMPE --extsize=300 -B -q 0.01 --nomodel --broad for ChIP-seq. The correlation between the replicates was calculated using deeptools(32) (version 3.5.0) multibigwigsummary based on the bedgraph output files from MACS2. The same approach was used, from a “macro” bam file pooling all the reads of all the ATAC-seq samples using Samtools v1.13, to generate a file with all islet accessible regions (“macropeaks” file). The correlations were visualized using the plotCorrelation command (--corMethod pearson). Replicated peaks were defined as peaks present in two or more of the samples per genotype.

Quantification of ATAC-seq reads in peaks was performed using featureCounts(33) v2.0.1. RNA-seq data from (6) was realigned and quantified using Salmon(34) v1.3.0 with parameters -p 20 -- validateMappings. Differential analysis of quantified peaks (ATAC-seq) and gene expression (RNA-seq) was performed with DESeq2(35) v1.28.0. The bedtools (36) v2.29.2 intersect tool was used to identify ATAC-seq peaks that intersect with ChIP-seq peaks and SLIC-CAGE TSS positions. HOMER(37) v4.9 package was used for gene annotation of peaks for integration with gene expression data and for transcription factor motifs analysis, with the functions annotatePeaks, parameters -go -genomeOntology and -gene and findMotifsGenome, parameters -size 200 and all the islet annotated ATAC-seq peaks (“macropeaks” file) as background, respectively. GREAT(38) v4.0.4 was used for the assignment of nearby genes to ATAC-seq annotations with DEseq padj values <0.1 and default settings: Proximal: 2kb upstream and downstream, plus Distal up to 100Kb.

DAVID(39) (v Dec. 2021) was used for gene ontology (GO) enrichment analysis with default parameters and REVIGO (25) was used to summarize and visualise enriched GO terms with Padj < 0.05.

### Splicing analysis with rMATS

For splicing analysis, previously generated high-throughput sequencing data of pancreatic islets from LKB1 knockout mice were used(6). FASTQ files were downloaded for LKB1βKO (n=4) and Control (n=5) islets from SRA-Explorer (available from project PRJEB7122). FASTQ raw sequence files were quality checked with FASTQC (http://www.bioinformatics.babraham.ac.uk/projects/fastqc). Low-quality reads and adapter sequences were trimmed and further processed to a single length of 52 bp with Trimmomatic(40), to fulfil the length requirements of downstream splicing analysis with rMATS(41) Trimmed FASTQ files were submitted to mapping and identification of transcriptome-wide splicing events with rMATS turbo v4.1.0(42). Mapping to the mouse genome assembly GRCm38.p6 was performed by STAR aligner(43). Identification of transcriptome-wide splicing events was performed using GENCODE GRCm38.p6 (release M25) GTF reference file as annotation. LKB1βKO pancreatic islets were compared with the control group using a cut-off splicing difference of 0.001. Analysis of rMATS output splicing files was performed using maser package (https://github.com/DiogoVeiga/maser). Candidate alternative splicing events (SE (skipped exon), A5SS (alternative 5’ splice site), A3SS (alternative 3’ splice site), MXE (mutually exclusive exon), or RI (Retained intron)) were defined considering reads that span junctions only (JC), as defined by rMATS. Further filtering was performed for those events with coverage >20, deltaPSI >0.1, and FDR < 0.1.

### Data analysis of the SLIC-CAGE

To map SLIC-CAGE paired-end sequencing data in mouse islets, we used the GENCODE assembly annotation version GRCm38.p6 and the STAR alignment tool(43) v2.4.2a. In the first step of our analysis, we converted read pairs into single-read BED file format by selecting the beginning positions of each read. In the next step, we used the bedtools (36) v2.29.2 merge function to convert read starting positions into clusters within a 20 nt window. We then removed single read clusters from further analysis and considered them as noise. For each cluster, we selected the position with maximum number of reads as the Transcription Start Site (TSS), using a threshold of a minimum of 2 CAGE reads per peak.

### Identification of pri-miRNA TSS

SLIC-CAGE peaks of mouse islets were further processed by removing clusters containing junction-reads as junctions near intragenic pri-miRNA TSS are not expected(44). To separate junction reads, we used the following bash command with samtools(45) v1.17: samtools view BAM | awk ’($6 !∼ /N/)’ | samtools view -bS -t ./GRCm38.p6.genome.fa.fai - > BAM.no_junction_reads.bam Each CAGE cluster was then intersected with ATAC-seq narrow peaks and ChIP-seq of histone marks H3K4me3 clusters as described above using bedtools (36) v2.29.2 intersect tool and allowing a 50 bps flanking region to broaden the area of the intersect. The first set of miRNAs was then classified relative to protein-coding genes by intersecting annotated miRNAs from GENCODE (vM25 annotation) with protein-coding gene promoter regions spanning 200 kbps upstream from the TSS position. For the intergenic miRNAs, we removed all GENOCODE-annotated promoter regions except for lncRNAs and annotated TSS within 200 kbps of miRNAs as potential miRNA TSS. The classified CAGE peaks were then merged into a final table of 6197 potential miRNA TSSs (see Results).

All the pipelines and python scripts for processing CAGE-seq data and pre-miRNA TSS identification used in this study are deposited and available on github (https://github.com/nebo56/miRNA_TSS_classification_using_CAGE-seq).

## RESULTS

### LKB1**β**KO islets display an altered chromatin landscape

To investigate a role for LKB1 in the control of the chromatin landscape of pancreatic β-cells, we compared the chromatin accessibility profiles of pancreatic islets from mice with β-cell-specific deletion of LKB1(6) (LKB1βKO) with that of littermate controls using ATAC-seq (see Methods). Analysis of fragment size distribution of these libraries (Supplemental Figure 1A) demonstrated good quality of the ATAC-seq data with clear nucleosome phasing characterized by an enrichment in fragments < 100 bp, corresponding to open chromatin and a typical peak of ∼200bp (mono-nucleosome regions), with other less abundant, larger peaks (multi-nucleosomes). Using MACS2 (46) for peak calling (q < 0.01), we identified 209,194 accessible chromatin regions, 67,612 of which were detected in at least two samples of either genotype (Supplemental Tables 1, 2). Correlation analysis confirmed a high degree of concordance between experimental replicates (Pearson coefficients > 0.93, Supplemental Figure 1B).

We next used the DESeq2 package(47) to perform differential accessibility analysis of the identified open chromatin regions between control and LKB1βKO islets. Principal component analysis (Figure 1A) revealed a marked effect of the ablation of β-cell LKB1 on the chromatin landscape of pancreatic islets. We identified 1,693 and 7,400 open chromatin regions (p-adj < 0.1, Figure 1B) with significantly reduced (less “open”) and increased (more “open”) accessibility, respectively (Supplemental table 3). GREAT analysis(38) assigned the genomic regions with reduced and increased accessibility to 1,437 and 6,491 genes, respectively (Figure 1C, Supplemental Table 4). Pearson correlation analysis identified a positive correlation between fold changes of differential accessibility (ATAC-seq) and differential RNA expression (RNA-seq), and this was most evident for peaks located in close vicinity (<1500 bp) to annotated TSS (Figure 1D). We have previously shown(6) that LKB1βKO islets display increased expression of dozens of neuronal genes and, consistent with this, gene ontology analysis of the genes associated with chromatin regions with differential accessibility revealed a strong enrichment in several neuronal pathways (Figure 2, Supplemental Table 5), including ‘nervous system development’ (p-adj < 0.001). Examples include the regulator of glutamate signalling and neuronal plasticity(48) *Nptx2* (neuronal pentraxin 2) and *Dlgap2* (discs, large(Drosophila) homologue-associated protein 2), a protein of the postsynaptic density scaffold involved in glutamate signalling (49).

**Figure 1.**
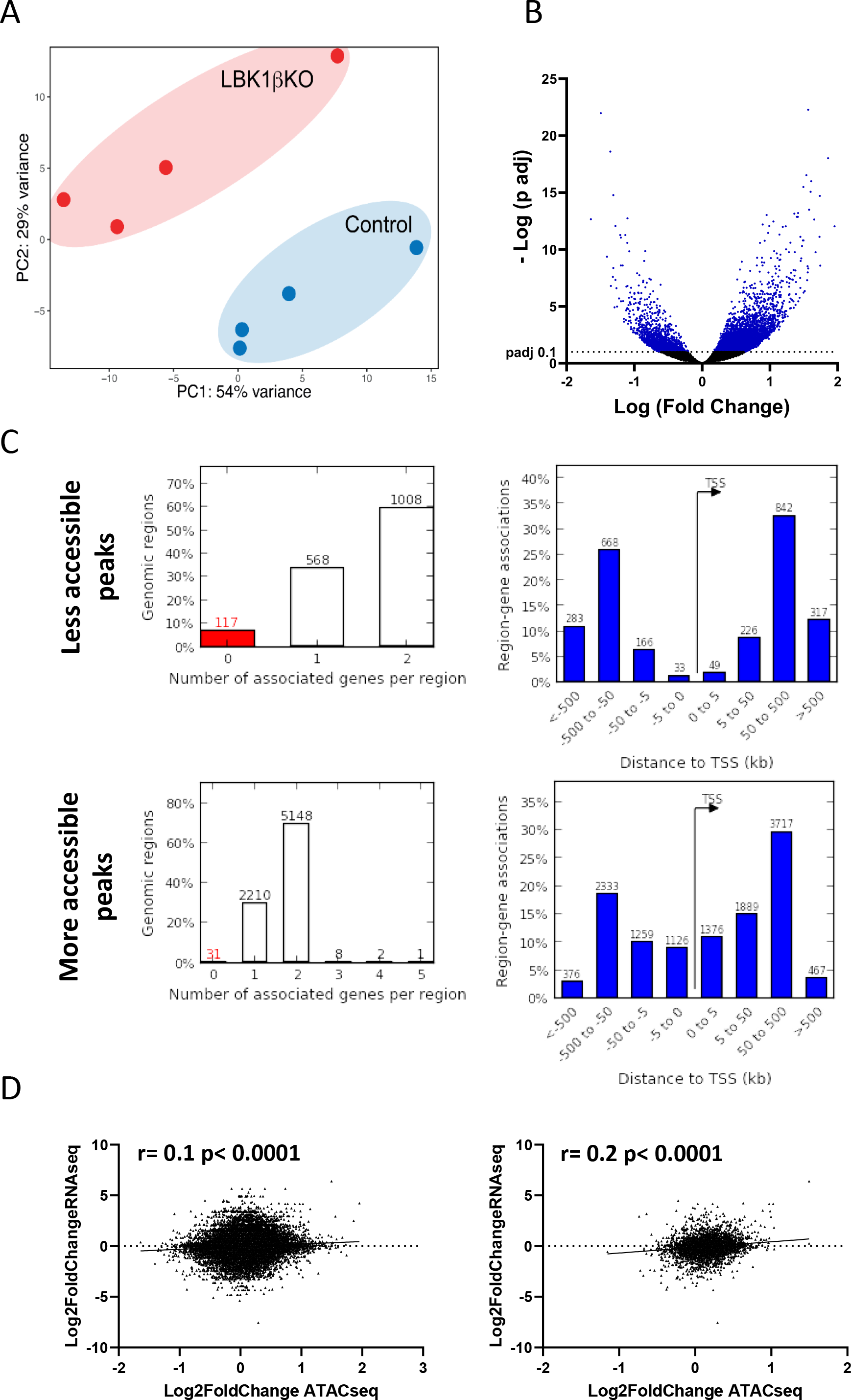
Elimination of LKB1 alters the chromatin landscape of pancreatic β-cells. A) PCA plot of ATAC-seq profiles of mouse islets with β-cell specific knockout of LKB1 (LKB1βKO) and littermate controls (n=4). **B)** Volcano plot showing chromatin regions with significantly different (padj < 0.1, in blue) accessibility LKB1βKO *versus* control islets. Each dot represents one ATAC-seq peak. **C)** Assignment of less (top) and more (bottom) accessible regions in LKB1βKO islets and TSS of their associated genes, predicted by GREAT(38) using default settings (see Methods). Left hand-side panels: number of associated genes per region; Right hand-side panels: distribution of distances between each differentially accessible region and associated TSS in kb. Numbers above each bar denote the respective gene count. The y-axis shows the fraction of all associated genes. **D)** Correlation analysis between log2 fold changes in chromatin accessibility and transcript expression in LKB1βKO islets. Pearson correlation values are shown. Accessible chromatin regions were assigned to expressed transcripts using HOMER. Right-hand side panel shows data on regions annotated within 1500 bp from the closest TSS.

**Figure 2.**
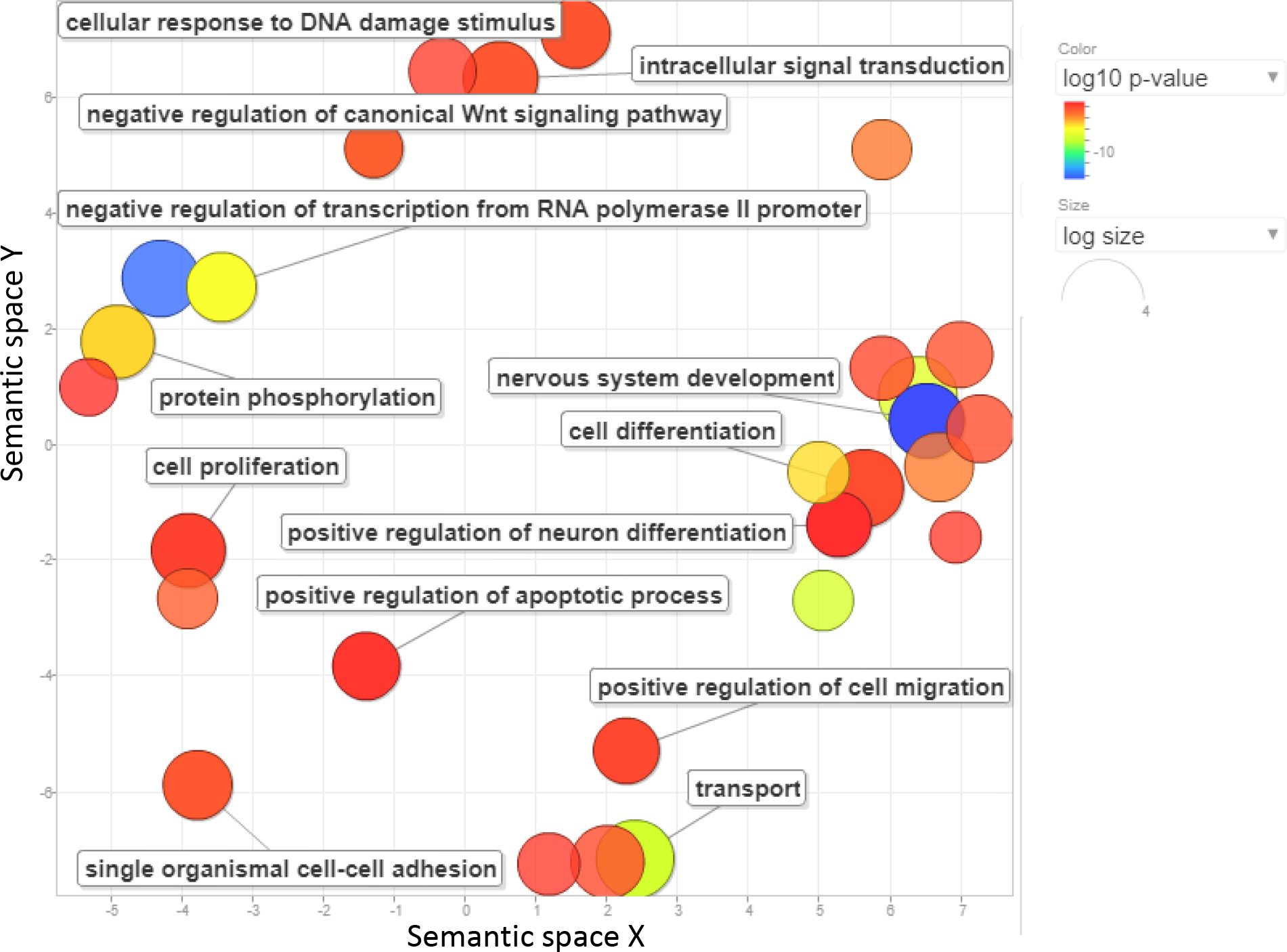
Gene ontology analysis of the genes associated with chromatin regions with differential accessibility in LKB1βKO. Gene ontology (GO) enrichment analysis for genes associated with differentially accessible regions was carried out with DAVID(39). REVIGO(105) was then used to summarize and cluster enriched GO terms (padj < 0.05) according to semantic similarities. The table view in the lower part of the figure is truncated (see Supplemental Table 5 for a full list); representative terms for each cluster are shown. Bubble colour (legend in upper right-hand corner) and size (the bigger the bubble, the smaller the value) indicates the padj-value.

We also observed enrichment in pathways related to gene transcription (i.e. ‘positive regulation of DNA-binding transcription factor activity’ (p-adj < 0.001), Figure 2, Supplemental Table 5), consistent with the profound effects on gene expression observed in LKB1βKO islets(6). Additionally, we detected the enrichment of several other biological pathways and functions that have been experimentally demonstrated to be altered in LKB1βKO animals by us and others (6, 7, 13). These included cell proliferation (p-adj = 0.002), cell/focal adhesion (p-adj < 0.001), calcium signalling (p-adj < 0.001) and insulin secretion (p-adj < 0.001). These results suggest that changes in chromatin accessibility resulting from LKB1 deletion are an important mechanism contributing to LKB1 action in pancreatic β-cells.

### LKB1 controls accessibility of enhancers enriched in islet transcription factor binding motifs

To further classify the differentially open chromatin regions identified in LKB1βKO islets, we first integrated a series of publicly available datasets of mouse islet chromatin immunoprecipitation followed by high-throughput sequencing (ChIP-seq) of histone modifications (29, 30) that are usually associated with active *cis*-regulatory elements, such as enhancers and promoters. Since the ambiguity of these marks makes it difficult to distinguish promoters from enhancers, we profiled active sites of transcription in mouse islets using Super-Low Input Carrier Cap analysis of gene expression (SLIC-CAGE). Thus, open chromatin regions enriched in H3K4me3 and containing SLIC-CAGE-determined TSS (a total of 15,477 peaks) were defined as active promoters in mouse islets, whilst those enriched in H3K27ac and/or H3K4me1 in the absence of H3K4me3 were defined as enhancers (41,452 regions). Accordingly, we annotated 52 ATAC peaks with reduced accessibility in LKB1βKO islets as active promoters while 419 corresponded to enhancers (Supplemental Table 6 A, B). Regarding the regions with increased accessibility, 2,857 corresponded to enhancers while 2,387 were classified as active islet promoters (Supplemental Table 7 A, B).

LKB1 is a protein kinase without reported DNA binding properties(50). Nevertheless, the enzyme has recently been shown to modulate chromatin accessibility and transcription factor (TF) expression during metastatic progression via signalling mediated by salt-inducible kinases (SIK) (20). Thus, we hypothesized that TF binding to chromatin regions with differential accessibility (control *versus* LKB1 null cells) contributes to the actions of LKB1 in maintaining chromatin accessibility in pancreatic islets. Indeed, transcription factor motif enrichment analysis within these regions(37) revealed that enhancers with increased accessibility were enriched for motifs corresponding to several key islet transcription factors involved in maintaining β-cell identity and often dysregulated in diabetes(51), such as FOXA1, FOXM1, FOXO2, MAFA, NEUROD1 and RFX6 (Figure 3, Supplemental table 8).

**Figure 3.**
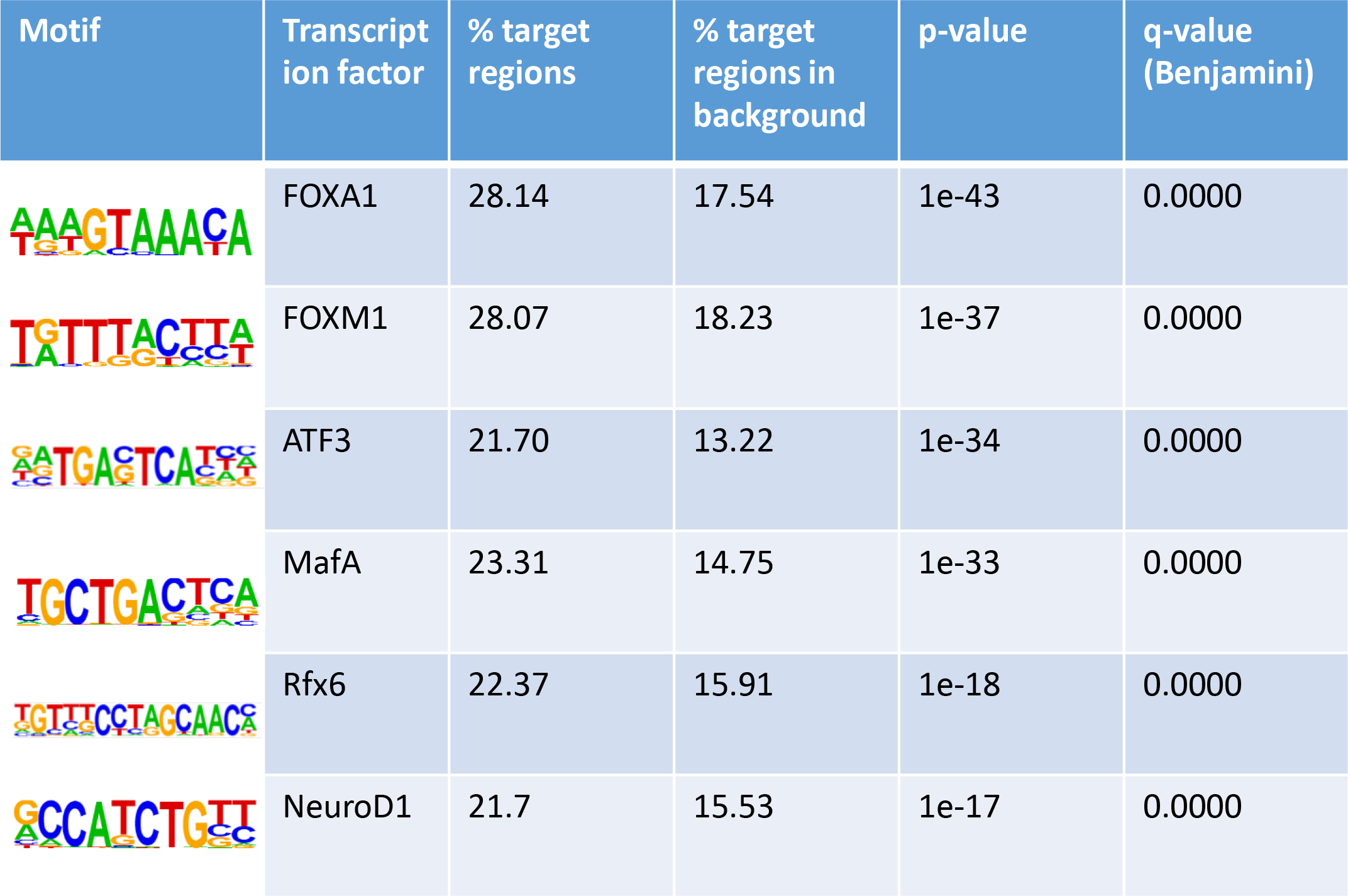
Transcription factor (TF) binding enrichment analysis of enhancer regions with increased accessibility in LKB1βKO islets performed with HOMER identifies several key islet transcription factors. Top 6 hits are shown, see Supplemental table 8 for a complete list.

Enhancer regions with decreased accessibility following LKB1 elimination displayed only a modest enrichment in TF binding motifs corresponding to LHX9/En1 and RORγ (*RORC*) (Supplemental Table 9). Whilst the function of LHX9/En1 in islets remains unclear, RORC has been previously shown to promote insulin secretion in a rat insulinoma cell line and is downregulated in islets from diabetic donors(52). On the other hand, active promoters with increased accessibility upon LKB1 elimination were enriched in motifs recognized by ubiquitous transcription factors such as Nuclear transcription factor -Y (NF-Y) and SP (specificity protein) family members (Supplemental Table 10) which often co-occupy mammalian promoters (53). These TF play essential roles in cell cycle, proliferation, and apoptosis (54, 55). Interestingly, we also found enrichment in NRF (Nuclear factor erythroid 2-related factor) binding motifs, a transcription factor involved in mitochondrial biogenesis and oxidative phosphorylation(56) and members of the ETS family, which can be activated by nutrients(57). Promoters with reduced accessibility did not show a significant enrichment for specific TFs, which is expected given the very small number of regions compared to the other sets of regions.

These results indicate that deletion of LKB1 in β-cells alters chromatin accessibility which results in, or reflects, altered access of islet-specific transcription factors to enhancers, and of ubiquitous transcription factors to promoters. This leads to concomitant changes in the expression of genes involved in processes important for the maintenance of β-cell identity and function, such as cAMP, AMPK and calcium signalling or neuronal genes.

It has been proposed that active enhancers are often transcribed producing enhancer RNAs (eRNAs), which are usually capped but non-polyadenylated (58). eRNAs play important roles in the regulation of gene expression for instance by altering chromatin structure(59)and stabilising enhancer-promoter loops(60) and hence may be involved in mediating some of the actions of LKB1 on the islet transcriptome. These transcripts cannot be detected using traditional RNA-seq experiments which usually involve mRNA enrichment via selection of polyadenylated mRNAs. In contrast, SLIC-CAGE detects any transcribed and capped transcript (27). Enhancer transcription tends to associate with regulatory activity(61) and, in islets, it could pinpoint genetic variants with functional relevance(62). To investigate a possible role for LKB1 in regulating the activity of transcribed enhancers, we scanned the differentially open enhancers for the presence of SLIC-CAGE-determined TSS. Only 211 and 15 enhancers with increased and reduced accessibility in LKB1βKO islets, respectively, contained SLIC-CAGE signals. These enhancers only showed a modest enrichment in transcription factor motifs, including the islet-specific MafA, FOXM1 and Rfx6 (Supplemental Table 11).

It has also been proposed that genes required for pluripotency and for the regulation of gene expression during development are enriched in “bivalent domains”, characterized by the simultaneous presence of H3K4me3 and H3K27me3(63). Others have suggested that the loss of silencing at bivalent promoters occurs in T2D human islets and in mouse islets after exposure to a high-fat-diet, contributing to the loss of β-cell differentiation (29). In healthy human islets, only a minority of genes carry both marks simultaneously, including those associated with neuronal transcription factors(64). Our previous work(6) demonstrated the loss of β-cell identity and increased expression in neuronal genes in LKB1βKO islets. Thus, we questioned whether elimination of LKB1 might affect chromatin accessibility of bivalent promoters. We identified 1,757 regions of open chromatin that contained both H3K4me3 and H3K27me3 and thus were classified as bivalent promoters. Only 127 and 14 of the regions with bivalent promoter marks were significantly more or less accessible, respectively, in LKB1βKO islets. Notably, gene ontology (GO) analysis of the 157 genes (Supplemental table 12) associated with these regions revealed a significant enrichment in several biological pathways related to neuronal function as well as cell to cell adhesion (Supplemental table 13). Nevertheless, given the small number of genes, it was not possible to detect significant enrichment of specific TF motifs in these regions.

### LKB1 has a minor effect on mRNA splicing

Given the widespread effect of LKB1 in gene expression(6), we wondered whether this kinase also plays a role in the regulation of splicing in islets. We thus used replicate multivariate analysis of transcript splicing (rMATS)(41) to compare the splicing patterns of LKB1βKO and control islets (65) detected by RNA-seq. The analysis identified 9,943 splicing events after filtering those annotations with a read coverage >20. Of these, 12 splicing events were significantly different in LKB1βKO islets (delta percent splice inclusion (dPIS)>0.1 and false discovery rate (FDR) < 0.1), albeit the extent of the differences were small (Supplemental Table 14). The three differentially regulated splicing events correspond to increased inclusion of a microexon in the genes *Papss2*, *Chchd3* and *Gas5* (Supplemental Figure 2). Of these, only *Gas5*, a lncRNA associated with the effects of glucocorticoids on insulin secretion, has been studied in the context of β-cells(66). Whether the small effect observed in the splicing of these genes has any impact on gene expression and β-cell function remains to be investigated. Overall, these results suggest that LKB1 has little to no role in splicing regulation.

### LKB1 regulates miRNA transcription

During the last two decades, miRNAs have emerged as essential regulators of virtually every biological process important for β-cell function (67). Islet miRNAs are regulated in response to stressors and dysregulation of miRNA expression occurs with and contributes to the development of diabetes(68, 69). Our group has previously identified 22 islet miRNAs regulated by AMPK and demonstrated that this enzyme mediates the glucose-dependent regulation of miR-184, a miRNA important for beta cell compensation during pregnancy and obesity (70–72), possibly at the level of transcription(23). Whereas miR-184 expression was concomitantly reduced in LKB1βKO islets(23), the effect of this kinase in the expression of other miRNAs, as well as the exact molecular mechanism by which LKB1/AMPK control miRNA expression was not interrogated.

MiRNAs are transcribed as longer primary transcripts (pri-miRNAs) that are rapidly processed into mature, functional, miRNAs (73). Due to this rapid processing, traditional RNA-seq and CAGE experiments have failed to annotate miRNA TSS in pancreatic islets up to now. We thus decided to take advantage of our highly-sensitive SLIC-CAGE and ATAC-seq data in islets, and of the published datasets of ChIP-seq of histone marks as described above, firstly to map miRNAs TSS specifically in mouse islets and, subsequently, interrogate the effect of LKB1 deletion on miRNA transcription.

To identify potential miRNA TSS, we first selected CAGE clusters overlapping with ATAC-seq and H3K4me3 peaks. Next, we employed a multi-step approach, similar to that previously described by Marsico et al (PROmiRNA(44)). The first set of miRNA TSSs was annotated relative to the promoter regions of protein-coding genes by intersecting CAGE peaks for all miRNAs that overlapped protein coding genes. We then excluded CAGE clusters from all promoter regions of coding genes, resulting in a set of potential miRNA TSS positions. To further refine this set, we selected only those promoters within 200kb of the annotated miRNAs, as previous research has shown that pri-miRNAs can extend hundreds of kb in length (see Methods for more details).

Using this approach, we identified 6,197 potential *MIRNA* TSS corresponding to 824 unique miRNAs (Supplemental Table 15). To identify miRNAs regulated by LKB1 at the transcriptional level, we examined which *MIRNA* TSS were located within chromatin regions with altered accessibility in LKB1βKO islets. We found four TSS within regions with lower accessibility, suggesting reduced transcription, in LKB1βKO islets. This included three newly identified miR-184 TSSs ∼120Kb upstream the annotated *Mir184* (Supplemental table 16). Additionally, 682 miRNA TSS, corresponding to 220 miRNA genes, were located at regions with higher chromatin accessibility (Supplemental table 16), suggesting increased expression. Notably, there was little overlap between these miRNAs and those altered by AMPK elimination as previously described using RT-qPCR: 63 miRNAs with TSS within regions with differential accessibility in LKB1βKO islets were present in the RT-qPCR panel we used in AMPKdKO islets in the past(23), but only 4 of these miRNAs (miR-184-3p, miR- 29a-5p, miR-378a-3p and miR-125b-5p) were dysregulated in both models.

Four miRNAs of those with TSS located within regions of increasing accessibility in LKB1βKO islets were of special interest, due to their important roles in β-cell function and identity. MiR-26a-5p (miR-26a) is up-regulated during islet postnatal differentiation, enhances endocrine cell differentiation(74), and modulates mature β-cell insulin secretion and replication(75). MiR-30d-5p (miR-30d) is a member of the miR-30 family important for the differentiation of pancreatic islet-derived mesenchymal cells into hormone-producing islet-like cells(76) and plays a protective role for mature β-cells(77). Additionally, miR-29b-3p (miR-29b) contributes to maintenance of β-cell identity by silencing the disallowed gene *Slc16a1* (Mct-1)(78) and regulates circadian genes expression in islets(79). Finally, high circulating levels of miR-125b correlate with hyperglycaemia in T1D and T2D patients (80, 81). Our recent work has identified miR-125b as a potent negative regulator of β-cell function which expression is regulated by glucose and AMPK/LKB1 in islets *in vivo* (82). Our results indicate that one or more of the TSS for these miRNAs are more accessible in LKB1βKO islets, suggesting more active transcription and potentially higher expression levels. To validate these findings, we performed RT-qPCR for the mature form of these miRNAs as well as miR-184, already known to be regulated by AMPK/LKB1, in control and LKB1βKO islets. As expected, LKB1βKO islets contained significantly lower levels of miR-184 and higher levels of miR-125b. Additionally, as hypothesized, we detected significantly higher levels of miR-26a and miR-30d in LKB1βKO islets (Figure 4). MiR-29b content was also higher in LKB1βKO islets though this difference did not reach statistical significance. Thus, our data support the suitability of our approach to map islet-specific miRNA TSS and identified β-cell miRNAs that are regulated by LKB1 at the transcriptional level in β-cells.

**Figure 4.**
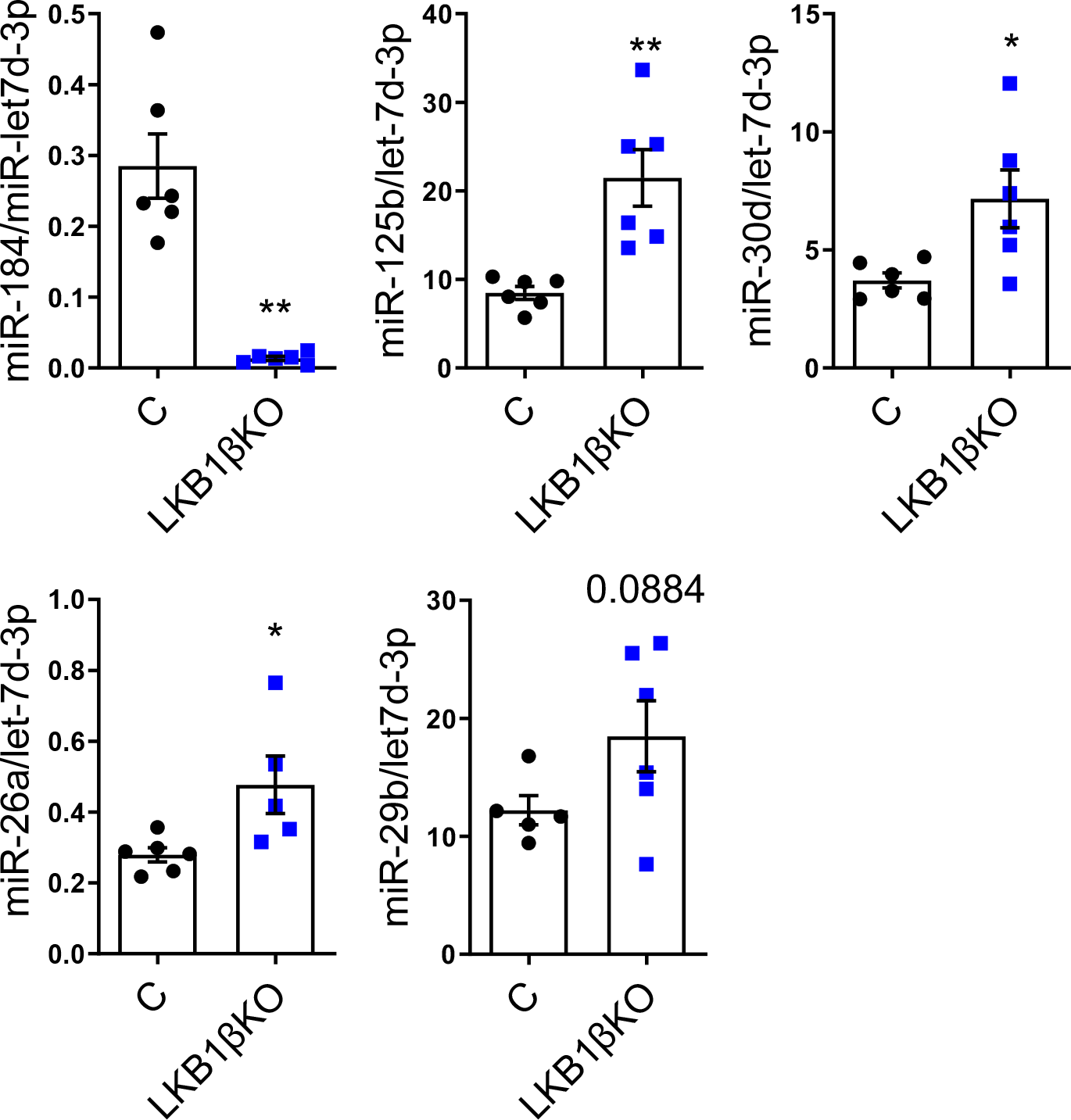
LKB1 regulates miRNA transcription. Expression of selected mature miRNAs was assessed by RT-qPCR in isolated islets from LKB1βKO (blue) and littermate control (C, black) mice. Each dot represents islets from a single mouse. Data are expressed as relative to the endogenous control let-7d-3p. *P < 0.05, ***P < 0.001, ****P < 0.0001 (Student’s t test).

Altogether, these results reveal an important role of LKB1 in regulating pancreatic islet identity programmes through not only the regulation of islet chromatin landscape but also of appropriate miRNA expression.

## DISCUSSION

Our results demonstrate that the elimination of LKB1 from mouse pancreatic β-cells results in widespread chromatin accessibility changes. We have previously identified an important role for LKB1 in maintaining β-cell identity(6) and that islets with β-cell specific deletion of this kinase are characterised by dysregulated expression of hepatic and neuronal genes, as well as genes involved in glutamate signalling. Gene ontology analysis of the genes close to regions of differential chromatin accessibility indicates that changes in the epigenetic landscape of these islets play a major role in the altered gene expression of genes involved in neuronal processes. A striking example is *Nptx2*, encoding neuronal pentraxin 2 and involved in glutamate signalling and neuronal plasticity(48), which is upregulated in LKB1βKO islets(6). LKB1-dependent chromatin changes were also associated with genes involved in cAMP and calcium signalling, as well as with insulin secretion. These epigenetic changes may therefore contribute to the enhanced capacity of beta cells to secrete insulin in response to external stimuli after LKB1 deletion(5–7, 13).

In a recent study, Pierce et al(20) uncovered an important role for LKB1 in the regulation of chromatin accessibility during metastatic progression of lung carcinomas. Notably, the authors identified a strong enrichment in SOX and FOXA motifs within regions with increased accessibility in adenocarcinoma cells following LKB1 deletion. Interestingly, most of the effects of LKB1 deletion in chromatin accessibility in these cells were mediated by salt-inducible family kinases (SIK), whilst AMPK inactivation had little to no effect. This is somewhat in line with our previous findings indicating that LKB1’s actions on β-cell gene expression are, at least in part, independent of the inhibitory effect on AMPK activity, implicating other, as-yet unidentified, downstream kinases(6). Mechanistically, highly metastatic tumours contained low levels of LKB1, high levels of SOX17 and displayed higher accessibility to genomic locations enriched in SOX17 binding sites(20). The authors demonstrated that restoring LKB1 expression sufficed to reduce *Sox17* mRNA levels, and that SOX17 is necessary and sufficient to maintain accessibility at genomic regions that contain SOX-binding sites in LKB1-defficient, metastatic cells(20). Of note, no changes in *SOX17* mRNA levels were observed in LKB1βKO islets. Moreover, sites of differential chromatin accessibility were not enriched in binding motifs for TFs of the SOX- family, suggesting a different underlying mechanism for LKB1 action in β- *versus* carcinoma cells.

Most chromatin regions with altered accessibility in LKB1βKO islets corresponded to enhancer regions. In line with the findings of Pierce et al(20), FOXA1 binding sites were enriched in enhancer peaks which displayed increased accessibility in LKB1βKO islets. The potential for LKB1 to modulate FOXA1/3 levels and activity post-translationally has already been demonstrated in lung adenocarcinoma cells(83). Interestingly, FOXA1/2 elimination results in functional defects in mature β-cells and in increased expression of genes involved in neuronal differentiation(84), similar to those observed in LKB1βKO islets (6). Further experimental validation is required to demonstrate whether FOXA TF mediate some of the effects of LKB1 in β-cell identity *via* transcriptional regulation of gene expression. Other TF motifs enriched in upregulated enhancers include ATF3, RFX6, MAFA and NEUROD1, all well-established regulators of β-cell identity and fate(85). Thus, our study suggests a mechanistic link between the actions of LKB1 and these transcription factors. Of these, only *MafA* mRNA expression was significantly dysregulated in LKB1βKO islets and future studies will be needed to confirm whether, and how, their activity is modulated by LKB1 and downstream kinases in these cells.

Albeit that very few (52) peaks with reduced accessibility in LKB1βKO islets were annotated as promoters, over 2,000 peaks with increased accessibility corresponded to promoters. These promoters were enriched for binding motifs to NF-Y, whose specific deletion in β-cells resulted in reduced proliferation, disturbances to the actin cytoskeleton and several molecular defects leading to impaired GSIS(86). A link between AMPK activity and NF-YA protein expression has previously been observed in tumours(87) and may make a similar contribution here. It is worth noting that binding motifs for NRF1 were also enriched at these sites. LKB1βKO islets displayed enhanced GSIS despite containing swollen and cristae- defective mitochondria, which were also less capable of producing ATP and NAD(P)H in response to glucose and showed defective respiration as determined by OCR(16). It is plausible that the increased activity of NRF1 at specific promoters contributes to these respiratory defects, since *Nrf1* elimination from murine β-cells increased mitochondrial respiration and resulted in mitochondrial swelling (56).

An important outcome of this study is the accurate mapping of TSS for mouse islet miRNAs. It has been widely demonstrated that miRNAs are important regulators of β-cell development and function(22, 88, 89) and that dysregulation of islet miRNA expression is associated and contribute to the development of diabetes(69). It has also been widely shown that islet miRNAs are regulated in response to stressors such as hyperglycaemia, hyperlipidaemia, exposure to cytokines and others (69). In contrast, little is known regarding the molecular mechanisms, including altered transcription, that lead to miRNA dysregulation in pancreatic β-cells. This is in large part due to the lack of accurate primary miRNA (pri-miRNA) annotations. MiRNAs are initially transcribed as longer pri-miRNAs that are speedily processed in a two-step-wise manner into mature, ∼22ntde, miRNAs(90). The rapid nature of this processing has rendered conventional RNA-seq experiments ineffective to identify miRNA TSS. In the past few years, researchers have combined histone modification and RNA polymerase II ChIP-seq as well as DNase-seq to computationally predict miRNA TSS(91). More recently, techniques aimed at experimentally identifying TSS, such as cap analysis of gene expression (CAGE) or global run-on sequencing (GRO-seq), have been applied to miRNA TSS annotation by consortiums annotating TSS in hundreds to thousands of cell types and tissues(91, 92). Nevertheless, islet isolation is a skilled and specialized protocol, and often isolated islets are not included amongst the cells/tissues analysed. Moreover, islets constitute only a ∼1% of the pancreas and therefore annotations made using the whole organ cannot readily be extrapolated to islet tissue. Importantly, it is often the case that different TSS are predicted but their use is highly cell type-specific, underlying the need of developing cell-specific annotations of miRNA TSS.

Using SLIC-CAGE with isolated mouse islets we have been able to capture low abundance capped mRNA including pri-miRNAs. Multiomics integration of SLIC-CAGE, ATAC-seq and histone modifications data has then allowed us to map precisely the TSS and promoters of the mouse islet miRNome. These findings may facilitate future mechanistic studies on the regulation of miRNA expression during β-cell development, nutrient responses and in disease. They may also inform the design of genetic models of miRNA gain/loss of function for functional studies.

In the present study, we have used the above information to identify miRNAs transcriptionally regulated by LKB1, defined as those with TSS located in areas of differential accessible chromatin following LKB1 knockout. Only two miRNAs were associated with promoters that displayed reduced accessibility after LKB1 deletion. One of these, miR-184, is an important regulator of β-cell compensatory response to pregnancy and obesity and we (23) and others (72) have shown its expression to be controlled by glucose in an LKB1 and AMPK-dependent manner, possibly at the transcriptional level(23). We have previously demonstrated that LKB1βKO islets contain ∼30 and ∼5 fold less mature miR-184 and pri-miR- 184, respectively, than control islets(23). The data presented here identified five separate miR-184 TSS in islets, defined as those with accessible chromatin, H3Km4me3 and SLIC-CAGE signal in the corresponding strand (negative strand for miR-184), at ∼70, 100 and 120 kb upstream miR-184. This is in contrast to earlier suggestions of a single miR-184 promoter located ∼78 kb upstream the annotated miR-184 stemloop(93). A strong decrease in accessibility in LKB1βKO islets (∼0.49 fold, padj=2.08E-05) was observed only for the promoter region surrounding the ∼120kb site, indicating that transcriptional changes *via* that region underlie LKB1-mediated regulation of miR-184 expression.

Further validating our annotations, RT-qPCR experiments confirmed that the expression of other mature miRNAs, namely miR-125b, miR-26a, miR-29b and miR-30d with pri-miRNA TSS mapped in promoters with differential accessibility in LKB1βKO islets, was also significantly changed in LKB1βKO islets. Although we focused our attention on miRNAs previously described to play functional roles in β-cells, our results suggest that LKB1 regulates the transcription of over 200 miRNAs. Notably, there was little overlap between these miRNAs and those dysregulated following AMPK elimination in islets. This result is barely surprising, given the only partial overlap existing between dysregulated protein-coding genes in AMPKdβKO and LKB1βKO(6) and further emphasize that these two proteins partially act independently of each other.

A single miRNA often targets dozens of genes making it likely that LKB1-regulated miRNAs are significant mediators of some of the roles of this protein in β-cells. For example, members of the miR-29 family could contribute to the effect of LKB1 on GSIS or in cellular identity(78, 94–96) and increased miR-26a might contribute to the effects of LKB1 elimination in β-cell proliferation(74) and secretion (75), whilst miR-184 reduction is likely to contribute to increased proliferation, similarly to what has been observed during compensatory responses to pregnancy and obesity(72).

Less is known about a possible role for LKB1-regulated miRNAs in mitochondrial morphology, respiratory capacity, and other functional features of this organelle. We have recently observed that miR-125b is regulated by glucose in an LKB1/AMPK-dependent manner and that this miRNA has a negative effect in pancreatic β-cell function acting at different molecular and cellular levels. While we identified several genes associated with mitochondrial function as miR-125b targets in β-cells(97–99), the functional relationship between this, and possibly other LKB1-dependent miRNAs, and mitochondrial homeostasis in the β-cell warrants future investigation.

MiR-125b is encoded by two independent genes, *MIR125B1* and *MIR125B2* and processed from several different primary miRNAs originated from different TSS(100–102). Our data identified CAGE peaks at fourteen promoter locations, ten corresponding to *MIR125B1* and four on *MIR125B2* (Supplemental Table 15). Of those, only the TSS located ∼2Kb upstream of the annotated pre-miR-125b-2 is within a promoter significantly more accessible in LKB1βKO islets, suggesting a major role of this locus and that specific TSS in miR-125b transcription in islets. This agrees with our previous findings that CRISPR-mediated deletion of *MIR125B2* but not *MIR125B1* sufficed to reduce mature miR-125b >80% in a human β-cell line (EndoCβ-H1)(82). In contrast, miR-125b is mainly transcribed from *MIR125B-1* in neighbouring acinar cells(100), emphasizing the importance of studying miRNA regulation in a cell-specific manner.

## Study limitations

The data presented here contributes to a better understanding of the molecular mechanisms underlying the role of LKB1 in β-cell function. Nonetheless, these results should be considered in the light of a few limitations. First, our results have been obtained using whole mouse islets in which β-cells comprise only ∼ 60-80 % of all cells within the islet ((1) and see Introduction) and, though our deletion strategy is highly selective and efficient in deleting LKB1 from these cells (6, 103), the remaining cells in this preparation remain unaffected and may thus have reduced our power to detect changes in the β-cell complement. Furthermore, mouse islets have a different architecture and, to some extent, altered glucose responsiveness and secretory dynamics *versus* human islets. Little is known regarding the function of LKB1 in human islets and thus the translatability of our results to humans is uncertain. We note that Varshney et al.(62) generated no-amplification non-tagging CAGE libraries for Illumina next-generation sequencers (nAnT-iCAGE) to identify TSS in human pancreatic islets, but did not map pri-miRNAs TSS. SLIC-CAGE increases the nAnT- iCAGE sensitivity over 1000-fold, without compromising signal quality, and could thus be used in the future to increase TSS annotation coverage and resolution when working with low amounts of human islets(27). It is also worth noting that the primary miRNAs annotated here might also act as lncRNAs with putative miRNA-independent functions.

Secondly, our study falls short of identifying the downstream kinases responsible for the described effects of LKB1 on chromatin remodelling. Third, we have previously observed a small, albeit significant, increase in β- to α-cell ratio in LKB1βKO islets(6) and thus we are unable to fully rule out the possibility that some of the changes in chromatin accessibility observed here represent this small change in cellular composition. Nevertheless, this is unlikely to be significant, since we observed no differences in accessibility to the β-cell specific *Ins2* promoter (not shown) and, contrary to what we would expect due to a higher β- to α-cell ratio, chromatin accessibility on the α-cell specific glucagon (*Gcg*) promoter was increased, probably reflecting β-cell dedifferentiation. Single-cell -omics analyses, including single-cell RNA- and ATAC-seq, might address this limitation as well as the potential complications resulting from the known heterogeneity of the β-cell population itself(104).

## Conclusion

In summary, our work demonstrates that LKB1 is important to maintain the epigenetic landscape of mouse β-cells and identifies transcription factors and miRNAs which may mediate some of the effects of LKB1 on gene expression and cellular function in the pancreatic islet.

## Acknowledgements

AMS was supported by a New Investigator Research Grant (MR/P023223/1) and a Project Grant (MR/X009912/1) by the Medical Research Council (MRC, UK). GR was supported by a Wellcome Trust Investigator Award (212625/Z/18/Z), UK MRC Programme grant (MR/R022259/1), Diabetes UK Project grant (BDA16/0005485), CRCHUM start-up funds, an Innovation Canada John R. Evans Leader Award (CFI 42649), JDRF (JDRF 4-SRA-2023-1182-S-N), and CIHR (CIHR-IRSC:0682002550) project grants. GR also acknowledge support from the NIH (R01 DK135268). IC was an Academy of Medical Sciences Springboard Fellow (SBF005\1050). The Scheme was supported by the British Heart Foundation, Diabetes UK, the Global Challenges Research Fund, the Government Department for Business, Energy and Industrial Strategy and the Wellcome Trust. IC is recipient of a Sir Henry Dale Fellowship jointly funded by the Wellcome Trust and the Royal Society (224662/Z/21/Z). DN is recipient of a PhD studentship by the British Heart Foundation (FS/4yPhD/F/20/34128). This work was also supported by Wellcome Trust (Joint-Investigator award 106955/Z/15/Z) to B.L., and core funding from Medical Research Council (award MC_UP_1102/1) awarded to B.L. Research support to LB was from the Spanish Ministry of Science and Innovation- Instituto de Salud Carlos III (ISCIII), which finances grant PI19/00468, cofinanced by “Fondo Europeo de Desarrollo Regional” (FEDER) and Centro de Investigación Biomédica en Red de Enfermedades Neurodegenerativas (CIBERNED). This research was also supported by Ramon y Cajal (RYC2018-024397-I) and IKERBASQUE (RF/2019/001) fellowships awarded to LB. Most of the data analyses presented here were conducted on the Imperial College Research Computing Service, DOI: 10.14469/hpc/2232

## Data Availability

The data that support the findings of this study will be openly available upon publication in Gene Expression Omnibus.

## Contributions

N.H. designed experiments and analysed data; R. C. and G. P. performed islet isolation and RT-qPCR experiments; N.C. and B.L generated SLIC-CAGE data; D. N. and H.M assisted with ATAC-seq data analysis; L.B. performed splicing data analysis; I. C. obtained ATAC-seq data, assisted with data analysis and contributed to the experimental design, and to write the paper. A.M-S., obtained and analysed data, designed research studies and wrote the paper. All authors read and approved the manuscript. I.C., G.A.R. and A.M-S. should be considered Senior authors.

## ABBREVIATIONS

LKB1: (STK11) Liver kinase B1 (Ser/Thr kinase 11)
AMPK: AMP-activated protein kinase
ATAC-seq: assay for transposase-accessible chromatin with high-throughput sequencing
SLIC-CAGE: super-low input carrier Cap analysis of gene expression
KO: knockout
CRISPR/Cas9: clustered regularly interspaced short palindromic repeats/Cas9
MiRNAs: MicroRNAs
TF: Transcription factor

---

**Supplemental Figure 1.**
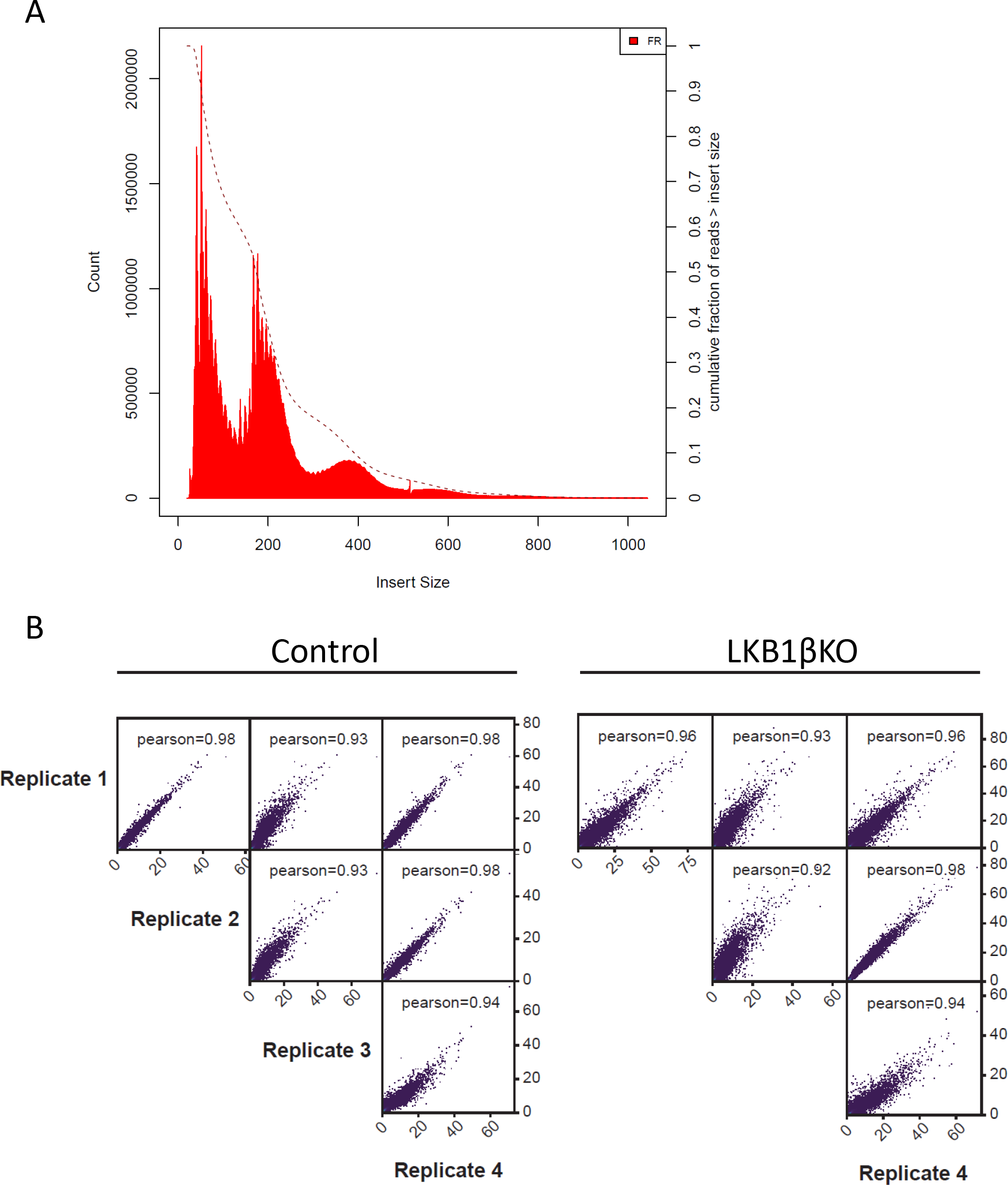
A) Fragment size distribution of ATAC-seq fragments obtained for all samples, as given by PicardTools (CollectInsertSizeMetrics). **B)** Correlation of genome-wide chromatin accessibility profiles (ATAC-seq) across experimental replicates.

**Supplemental Figure 2.**
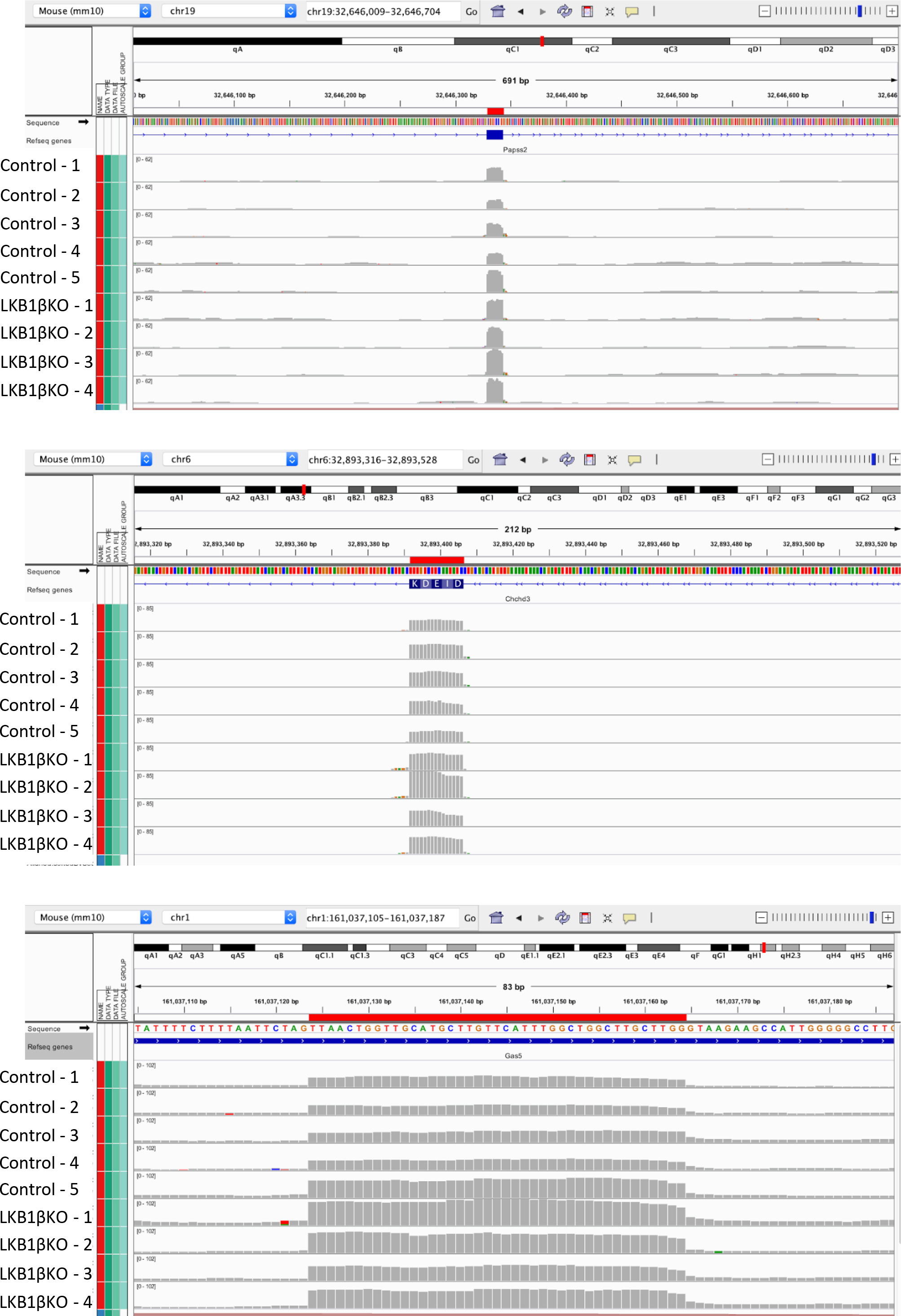
IGV browser visualisation of read coverage for microexons in genes *Papss2*, *Chchd3* and *Gas5*, showing significantly increased inclusion in LKB1βKO islets (4 bottom tracks) versus controls (5 top tracks).

